# Small Molecule Induced Toxic Human-IAPP Species Characterized by NMR

**DOI:** 10.1101/853549

**Authors:** Sarah J. Cox, Diana C. Rodriguez Camargo, Young-Ho Lee, Magdalena I. Ivanova, Vediappen Padmini, Bernd Reif, Ayyalusamy Ramamoorthy

## Abstract

In this study, the effect of CurDAc, a water-soluble curcumin derivative, on the formation and stability of amyloid fibers is revealed. CurDAc interaction with amyloid is structurally selective which is reflected in a strong interference with hIAPP aggregation, while showing weaker interactions with human-calcitonin and Amyloid-β_1-40_ in comparison. Remarkably, CurDAc also exhibited potent fiber disaggregation for hIAPP generating a toxic oligomeric species.

---

Small molecules have been sought out to interact with many different points in the aggregation process and can function as inhibitors, promoters, or disaggregators^[1]^. These tool compounds are advantageous for the targeting of many types of biological molecules, including amyloids. They are highly tuneable for specific functions, can bind tightly to the target protein, and can pass through cell membranes with little difficulty^[2]^. CurDAc, a curcumin derivative, has increased water-solubility and stability compared to its parent molecule (Figure S1). It has- been shown to inhibit hIAPP, both in the presence and absence of lipid bilayers, and negate hIAPP’s aggregation-induced lipid membrane disruption ^[3]^. In this study, the effects of CurDAc on three different amyloid peptides, amyloid-β (Aβ_1-40_), human islet amyloid polypeptide (hIAPP), and human calcitonin (hCT), were examined. While these three peptides have similar length ranging from 32-42 amino acids, they differ in their isoelectric point, hydrophobicity, and aggregation kinetics (Figure S2). These similarities and differences make them ideal for understanding the structural specificity of small molecules.

Fiber growth was monitored by utilizing Thioflavin-T (ThT),^[4]^ which is a widely used fluorescence reporter for amyloid fiber formation that gives kinetic and mechanistic information about amyloid aggregation (Figure 1). Using ThT fluorescence assays, both inhibition and disaggregation profiles were obtained for all three peptides, hIAPP (Fig.1 A&D), Aβ_1-40_ (Fig. 1 B&E), and hCT (Fig. 1 C&F). While CurDAc was able to modulate the aggregation of the three amyloid peptides, it displayed a distinct difference in potency. For hIAPP, as the concentration of CurDAc was increased from 0.06 to 2 molar equivalents, the overall final ThT signal intensity with respect to the control decreased indicating a decrease of fiber formation (Figure 1A). Not only is the overall intensity lower, but also the slope of the elongation phase is extended, indicating a possible change in the secondary nucleation step. CurDAc acted as a fibrillation inhibitor for hCT as well; however, it was not as effective as compared to hIAPP. For Aβ_1-40_, the concentration range that was tested on hIAPP and hCT had no effect on the inhibition of fibril formation (Figure 1C). Superstoichometric ratios (10-15 molar equivalents) were needed in order to see a depression of fibril formation in Aβ_1-40_ (Figure 1C). Interestingly, the lag-time at lower equivalencies of CurDAc was decreased by about 200 minutes, an effect that was not observed with the other two peptides. We believe that the change in kinetics based on the concentration of CurDAc is due to a colloidal inhibition effect and charge-charge repulsion of CurDAc with Aβ_1-40_^[5]^.

**Figure 1.**
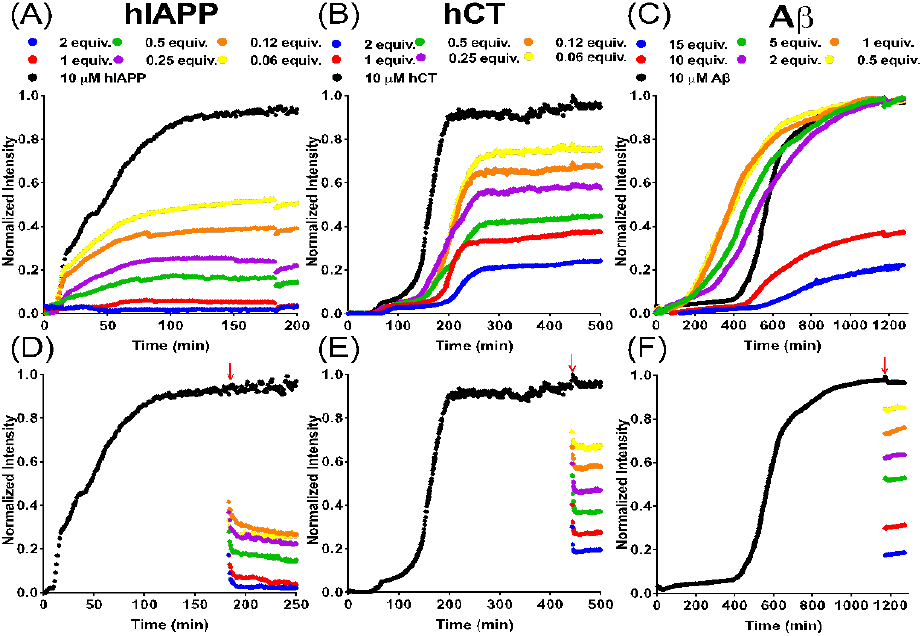
Inhibition and disaggregation of amyloid peptides by CurDAc. ThT fluorescence inhibition curves of hIAPP, hCT and Aβ_1-1-40_ (A-C respectively) in the presence of varying molar equivalents of CurDAc. Disaggregation curves of hIAPP, hCT and Aβ_1-40_ (D-F respectively) using the same molar equivalents as the inhibition experiments. Red arrow indicates when CurDAc was added.

Amyloid disaggregation as a method of removing plaque loads could be an avenue to reduced toxicity since plaques have been hypothesized to serve as nucleating sites for toxic oligomers ^[6]^. In Figure 1D, ThT curves show the disaggregation of hIAPP fibers upon the addition of CurDAc (red arrows) in the same stoichiometric ratios of CurDAc as in the inhibition experiments. A similar trend was seen for hCT in that there is an immediate drop in ThT intensity followed by a decline and plateau, however the steepness of decline is greater and the curve plateaus more quickly as compared to that observed for hIAPP (Figure 1E). For Aβ, we unexpectedly observed drops in intensity for all of the equivalents examined, however these curves are stable in intensity after the initial drop (Figure 1F). The initial steep decline in ThT intensity is followed by an eventual plateau. To rule out the possibility of ThT displacement we repeated these same experiments with higher excess of ThT and the same trend was observed (Figure S3), leading us to believe that another mechanism is operative. Using Isothermal Titration Calorimetry (ITC), the binding of CurDAc to Aβ_1-40_ or hIAPP was studied to further investigate the nature of the interaction (Figure S4 and Table S1). In the ITC experiments for hIAPP, no interaction was observed between hIAPP monomers and CurDAc. CurDAc interaction with hIAPP fibers was found to have a dissociation constant (K_d_) value of ~7 μM. For Aβ_1-40_ fibers, no binding could be detected for 10 molar equivalent concentrations of CurDAc. These data further support that the disaggregation effect is not a result of ThT displacement given the very similar binding constants of CurDAc to hIAPP fibers (~7μM) and ThT to amyloid fibers (~10 μM) ^[7]^.

Building on the ThT data of selectivity and potent disaggregation of hIAPP, we performed NMR experiments to obtain deeper understanding of this interaction. By observing the total proton NMR signal intensity of hIAPP in solution we analyzed the aggregation kinetics of hIAPP monomers in the presence of CurDAc (Figure S5). Interestingly, hIAPP in the presence of CurDAc at neutral pH loses signal intensity faster than in the absence of CurDAc. Most of the peaks in the NMR spectrum disappeared within 10 hours for the peptide alone. In presence of CurDAc, the peaks in the NMR spectrum disappeared in less than one hour, indicating the formation of NMR invisible species, such as a large off-pathway oligomer ^[8]^.

Human-IAPP is an unstable peptide under *in vitro* conditions, which is in accordance with quick self-assembly of the peptide to form aggregates under neutral conditions ^[9]^. It is also known that the formation of the N-terminus disulfide bridge is important for its aggregation kinetics; and in this NMR study, the oxidized form of hIAPP is used throughout. Due to the accelerated aggregation and signal loss at neutral pH, we lowered the sample pH to 5.3, which is the physiological pH of hIAPP when stored in insulin granules at high concentrations in the pancreatic beta cells ^[10]^. This strategy allowed us to perform longer NMR experiments on hIAPP and enabled us to complete additional analyses of the interaction with CurDAc due to the increased lifetime of the monomeric sample. At this pH, hIAPP remained relatively soluble over the course of 30 hours with only 20% signal loss (Figures S5 and S6). At a 1:1 molar ratio with CurDAc, the hIAPP signal decreased more rapidly with 70% of the signal lost after 20 hours and with a complete signal loss at 40 hours (Figure S5). This observation suggests that CurDAc is able to quickly initiate hIAPP aggregation and inhibits the fibrillar structure morphology of the aggregate species in the same way as other flavanols and confirms our initial observations using other biophysical methods. Using two-dimensional NMR techniques, such as SOFAST-HMQC, it is possible to obtain residue specific information of the CurDAc-hIAPP complex interaction (Figure 2A). In comparison to the reference ^1^H/^15^N HMQC spectrum of hIAPP alone, small CSPs were observed, in CurDAc containing samples, for residues in the N terminal and central regions as indicated by the peaks above the (horizontal) dashed line indicating the mean (Figure 2B). Most strongly affected residues include T4, C7, A8, Q10-N14, L16-S19, I26, and S29 and are highlighted on the monomeric structure (Figure 2B)^[11]^.

**Figure 2.**
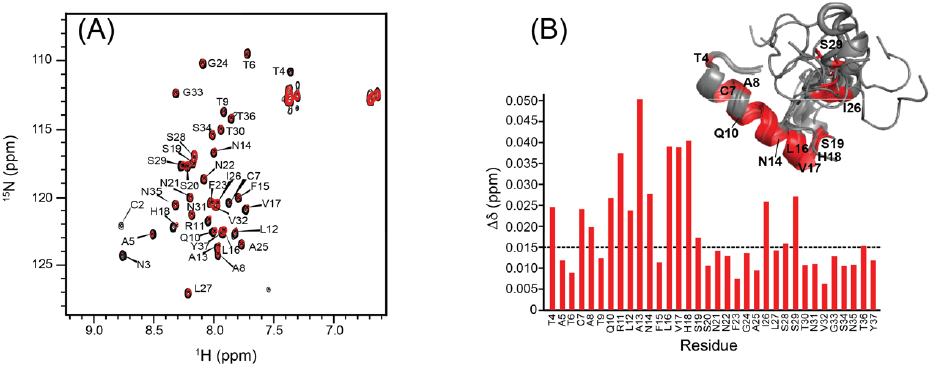
hIAPP monomer and CurDAc interactions characterized by NMR. (A) 2D ^1^H/^15^N SOFAST-HMQC spectra exhibiting the interaction between 100 μM hIAPP (black) with 100 μM CurDAc (red). (B) CSPs for the backbone amide ^15^N and proton resonances as a function of the amino acid sequence position. The dashed line indicates the mean for the observed CSPs with the stronger shift changes above 0.015 highlighted on the monomer structure of hIAPP (PDB:5MGQ).

Fibers represent a large insoluble form that is invisible for the traditional solution NMR experiments due to the presence (or incomplete averaging) of line-broadening anisotropic interactions such as dipolar couplings and chemical shift anisotropy^[12]^. In the event that portions of the peptides that form the fibers start to become soluble and are able to tumble sufficiently fast in the solution, the peptide signal will become narrow and the newly generated soluble peptide species can be identified and quantified. The starting NMR spectrum of the hIAPP fibers can be seen in Figure 3A (in black), in which very few peaks were observed. 72 hours after the addition of CurDAc, strong peaks appeared in the spectrum as shown in the amide region of the NMR spectrum (red). As shown in Figure 3B, after the addition of CurDAc to fibers, the signal intensity for the peaks observed in the proton NMR spectrum increased steadily over the course of 72 hours.

**Figure 3.**
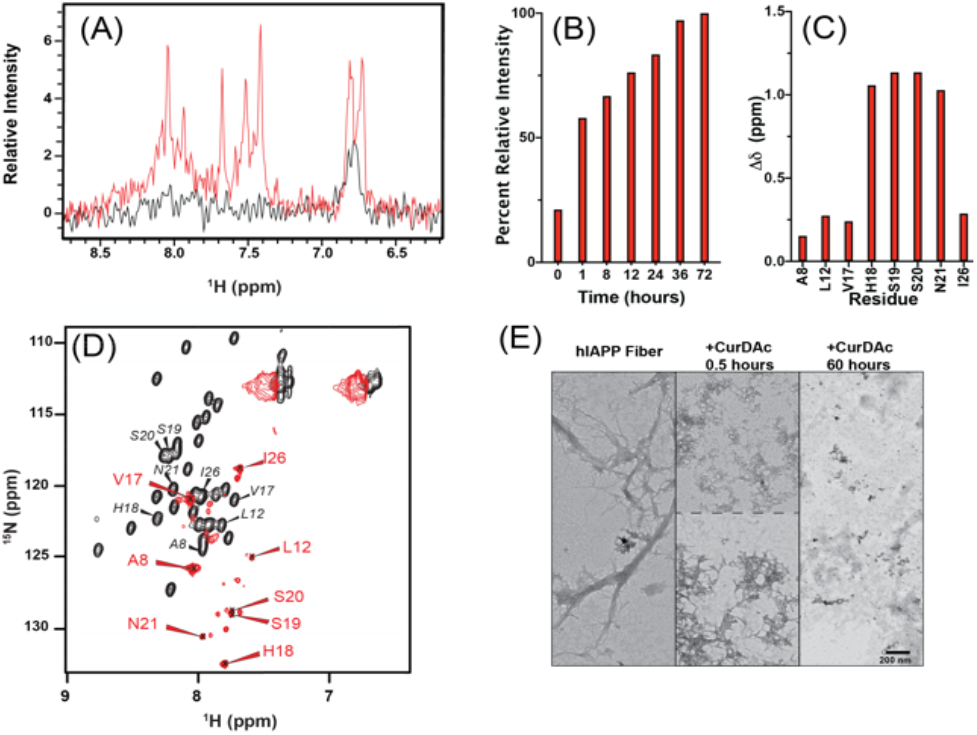
NMR observation of disaggregation of hIAPP fibers. (A) Proton NMR spectra of 90 μM hIAPP fibers (black) and 90 μM hIAPP after 72 hour incubation with CurDAc (red) at pH 7.4 in phosphate buffer. (B) 2D ^1^H/^15^ N SOFAST-HMQC NMR spectra of hIAPP monomers (black) and the disaggregated species (red) at a 1:1 peptide:CurDAc molar ratio. Peak assignments for the monomer are also shown for reference. (C) Normalized overall proton peak intensity of hIAPP fibers as a function of disaggregation by CurDAc over 72 hours. (D) Chemical shift perturbations for the backbone amide ^15^ N and proton resonances of the identified residues in reference to that of monomers. (E) TEM images of hIAPP fibers (left) and hIAPP species after 0.5 hours (middle) and 60 hours (right) disaggregation.

By using 2D SOFAST-HMQC NMR experiments, we were able to identify some of the peaks which appeared after 72h of adding CurDAc to hIAPP fibers. These resonances are attributed to mobile regions of the peptide and the disrupted aggregates of hIAPP fibers in solution (Figure 3D). 3D HNCA and 3D HNCOCA experiments were performed to identify some of these residues that appeared in the spectra of CurDAc-induced disaggregated hIAPP species (Figures S7 and S8). The 2D and 3D NMR spectra were not straightforward in their characterization due to the low concentration of disaggregated species which contributes to the observed NMR signal. However, these experiments do allow us to determine some of the residues and regions of hIAPP that have become visible as a result of disaggregation. From our partial assignment, we have identified the affected residues to be A8, L12, V17, H18, S19, S20, N21, and I26. All of these residues were also identified as residues affected by CurDAc interactions with hIAPP monomers by CSPs. Figure S9 shows the model of the amino acid sequence with the highlighted residues in red being the ones identified by the disaggregation of fibers. Out of the identified residues, V17-N21 are sequentially grouped, while A8, L12 and I26 are distributed randomly. While we were able to assign most of the visible resonances, a few resonances remained unassigned. It is possible that these unassigned peaks may belong to other residues that are near the three randomly distributed residues (A8, L12, I26).

When comparing the visible residues to known hIAPP monomer and fiber structures and models, residues V17-I26 can all be found in a loop region of the peptide. Due to this we believe that there are two possibilities for the absence of signals from other residues. One, possibility is that the residues are in a highly flexible/mobile loop region of an oligomeric structure, while unseen residues are buried within the oligomer causing NMR signal loss. The second possibility is that while a smaller species is becoming disaggregated, they are exchanging between bound and unbound states from the larger fiber structure. This could cause line-broadening of resonances corresponding to the residues that were not identified in the spectrum, making only these highly mobile loop residues that are not in exchange to be visible. Figure 3C shows the chemical shift perturbations of the identified residues. 6 out of the 8 residues have a significant upfield shift in the proton dimension and a downfield shift in the ^15^N dimension indicating a drastically different chemical environment as compared to that of the monomer. These chemical shift values do not fall in the typical range of known secondary structural chemical shifts for these residues. The CurDAc induced disaggregation of hIAPP fibers was corroborated using TEM (Figure 3E). TEM shows the difference of hIAPP fiber morphology before and after the addition of CurDAc. The disruption of aggregates increases over time as seen with decreasing amount and size of hIAPP fibers indicating the fiber disaggregation.

Given this exciting NMR data, we turned to other biophysical techniques to support our findings. By recording CD spectra of hIAPP over time after the addition of CurDAc, the secondary structural changes associated with the disaggregation of fibers were monitored. Before the addition of CurDAc, a strong peak was observed at 218 nm indicating a beta-sheet conformation (Figure S10). There was also a shoulder at 200 nm, which could indicate a portion of the peptide remaining in random-coil. Using CD fitting software, BeStSel, we analyzed the data to understand the possible distribution of conformations in the samples (Table S2). While a majority of the structural content was determined to be a beta-sheet, after the addition of CurDAc, a small helical conformation was observed between 30 minutes and 40 hours of the ongoing fiber dissociation. For Aβ_1-40_ and hCT, CD experiments indicated the formation of a stable beta-sheet secondary structure in the same time span as the ThT curves which was maintained after the addition of CurDAc (Figure S11). The CD samples were examined using TEM, and fibers were observed for all three different peptides before the addition of CurDAc. Since CurDAc’s effect on hIAPP is most pronounced, we monitored TEM over an extended period of time in accordance with NMR and CD experiments (Figure S10) in which the remodelling observed in the above-mentioned experiments can be visualized. For hCT after the addition of CurDAc, more amorphous type aggregates can be seen after remodelling with two molar equivalents (Figure S12). For Aβ_1-40_, the fibers maintained their architecture at 10 molar equivalents of CurDAc but appeared to be overall shorter in length than before addition (Figure S12).

There is considerable interest in understanding cellular toxicity of amyloid species in order to evaluate the potency of small molecule amyloid inhibitors and potentially to develop compounds to treat the related amyloid disease. While fibers are thought to be relatively inert, cytotoxic species are pointed at pre-fibrillar oligomers in the case of many different amyloid related diseases. In this study, the MTT dye reduction assay and CellTox green assays were utilized. (Figure 4). Freshly prepared monomers, or fibers grown for 24 hours, were added to the rat pancreatic islet cell line, RIN-5F, at varying concentrations and were allowed to incubate with adherent cells for 24 hours before measuring toxicity. It was found that the fibers exhibited greater toxicity than the monomers by decreasing the cell viability by greater than 40 percent at the highest concentrations used. When monomers and fibers were treated with CurDAc after 24 hours, the toxicities of both were increased in comparison to the non-treated samples. It is interesting to note that the toxicities of both monomers and fibers treated with CurDAc, exhibited very similar levels of toxicity indicating a level of similarity between the two. The toxicity of CurDAc to its parent compound curcumin was also compared, and it was found that both were nontoxic by the MTT assay (Figure S13). In order to corroborate our findings using the MTT assay the CellTox Green assay was utilized (Figure S14). This assay was in agreement with MTT results, in which we observed increased toxicity for the peptide that had been treated with CurDAc after 13 hours of incubation. These results indicate co-incubation of hIAPP monomers or fibers with CurDAc promotes the formation of toxic species. Statistical significance calculations were performed using ordinary one-way ANOVA tests with Tukey’s multiple comparisons. All values of the comparisons are shown in Table S3.

**Figure 4.**
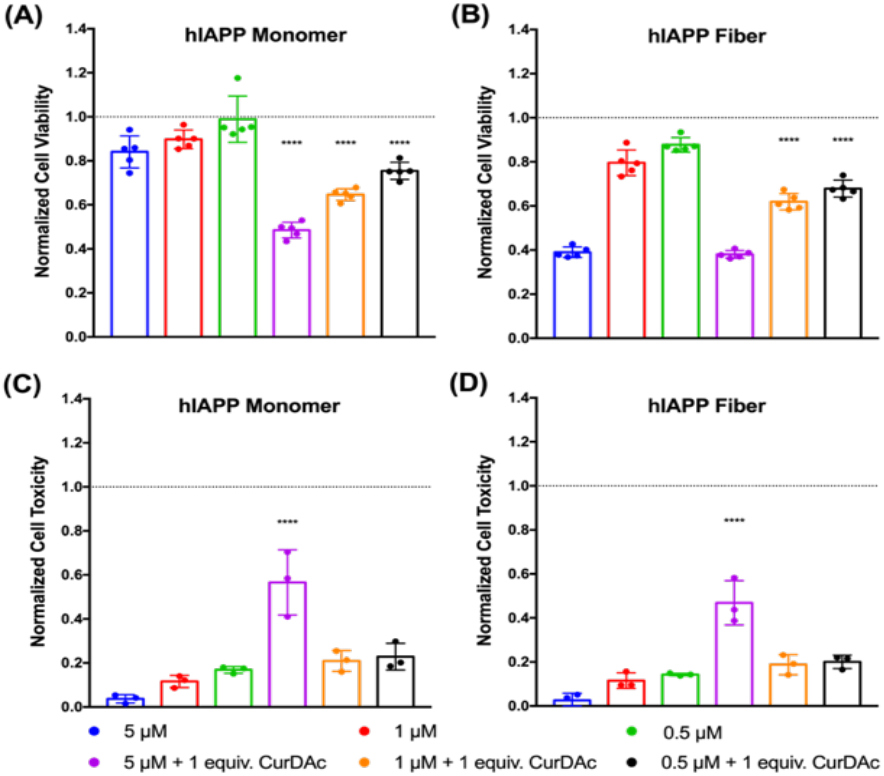
Cell toxicity measurements of hIAPP monomer and fibers in the presence and absence of CurDAc. RIN-5F cell viability measured by MTT reduction in the presence of hIAPP monomers and fibers without CurDAc and after 24-hour incubation with CurDAc (A-B). Cell toxicity final intensity after 13 hours of monitoring by CellTox Green fluorescence (C-D).

While amyloid aggregation has been extensively studied, use of small molecules as a chemical biology tool provides advantages to traditional methods of stabilization. This study has shown the potential of CurDAc to act as a peptide specific chemical probe to study hIAPP and to disrupt fully mature fibers and prevent monomers from forming fibers. This is the first example in the literature of using NMR to follow fiber disruption at atomic resolution which allowed us to identify solvent exposed residues in the species formed after disaggregation. Most of these residues, which are in the loop regions of known monomer and fiber structures, have distinct chemical shift values. This suggests that these residues are in a new chemical environment and/or structurally distinct and may be part of a larger oligomeric species that is not visible for solution NMR studies. TEM served as a visual confirmation in which the fibers can be seen being broken down upon incubation with equimolar amounts of CurDAc. Notably, these disaggregated species were more toxic than the fibers or monomers to cultured cells. We believe that the results reported in this study can be used to provide a starting point to develop other small molecules for the purpose of amyloid modulation or oligomer characterization. Using tool compounds like CurDAc could provide new avenues to probe and isolate the pathways of oligomer formation, which may lead to further understanding of the toxic oligomer hypothesis.

## Conflicts of interest

There are no conflicts to declare.

## Acknowledgments

This study was supported by the NIH (AG048934 to A.R.)).

