## Supporting information for "Small Molecule Induced Toxic Human-IAPP Species Characterized by NMR"

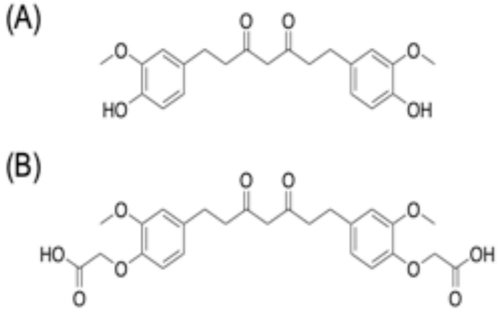


Figure S1. Chemical structures of curcumin (A) and its water-soluble derivative, CurDAc (B).

```
         1         10         20         30         40
   Aβ: DAEFRHDSGYEVHHQKLVFFAEDVGSNKGAIIGLMVGGVV
hIAPP: KCNTATCATQRLANFLVHSSNNFGAILSSTNVGSNTY
  hCT: CGNLSTCMLGTYTQDFNKFHTFPQTAIGVGAP
```

Figure S2. Amino acid sequences of the three different amyloid peptides (amyloid-beta, human-IAPP and human-calcitonin) used in this study. Color indicates charge of residue at pH 7.4: red is acidic, blue is basic, black is uncharged.

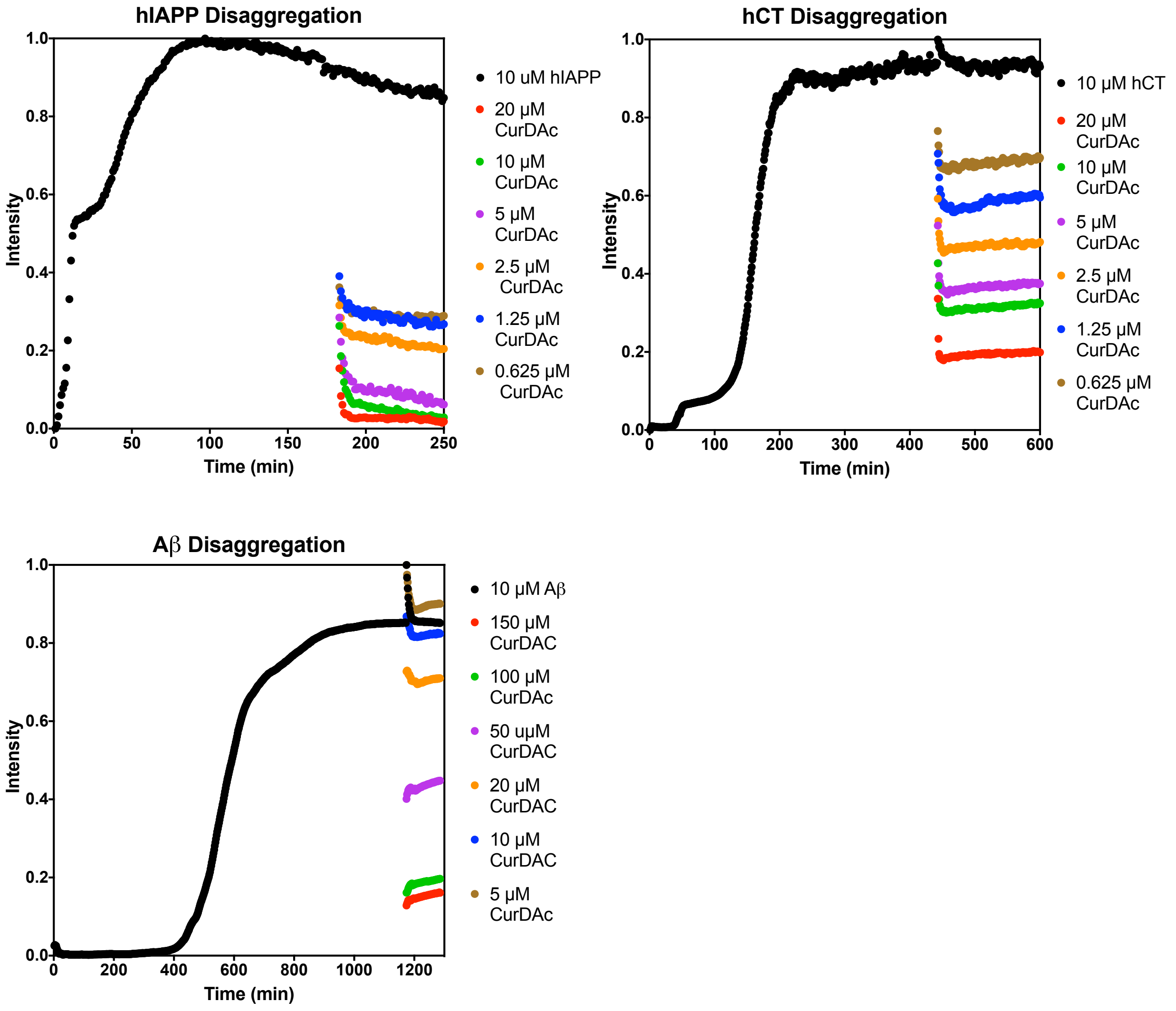


Figure S3. Kinetic disaggregation curves using 4 equivalents of ThT. 10 μM peptides (hIAPP, hCT and Aβ) were allowed to aggregate to form amyloid fibers, as shown in the black curve in each figure, with 40 μM ThT to monitor aggregation. Wells were then treated with the same stoichiometric ratios of CurDAc as shown in the previous experiments (Figure 1).

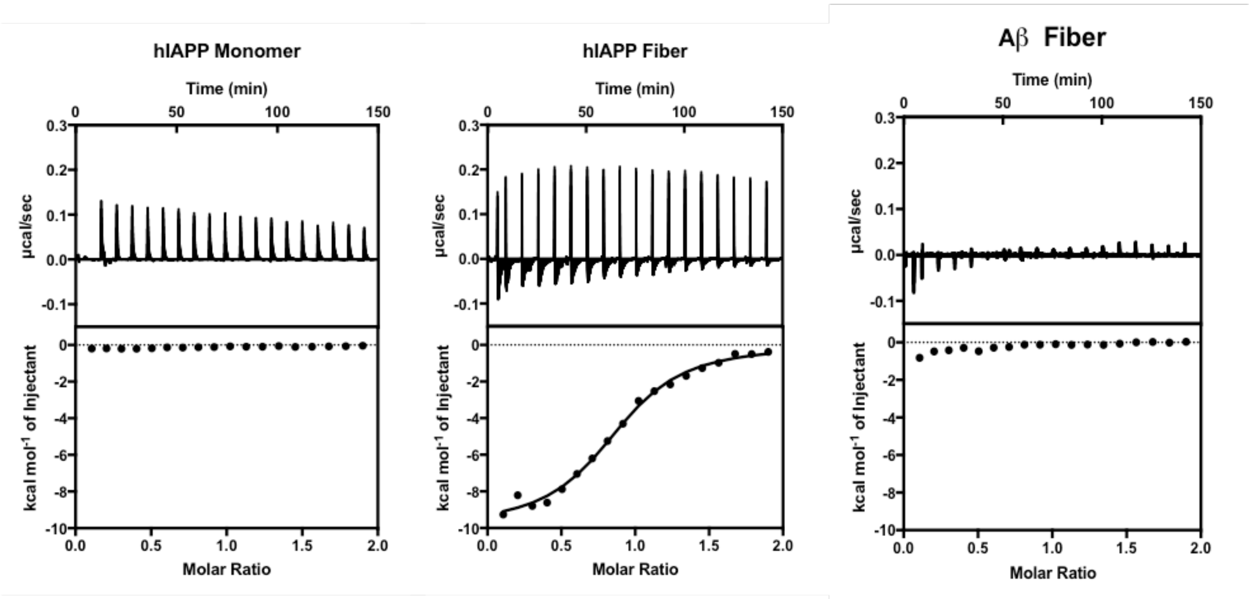


Figure S4. Isothermal titration calorimetry (ITC) experimental traces showing the monomer and binding with CurDAc over time. hIAPP monomer and Aβ fiber show no thermodynamic binding. hIAPP fibers with CurDAc show endothermic interaction.

| hIAPP Fibers + CurDAc | |
|---|---|
| **$\Delta G_{bind}$ (kcal mol$^{-1}$)** | -7.3 |
| **$\Delta H_{bind}$ (kcal mol$^{-1}$)** | -9.9 |
| **$-T\Delta S_{bind}$ (kcal mol$^{-1}$)** | 2.6 |
| **$K_d$ (μM)** | 6.8 |
| **n (value)** | 0.88 |

Table S1. hIAPP fiber and CurDAc parameters as determined from isothermal titration calorimetry (ITC) data (Figure S10). Dissociation constant ($K_d$), $\Delta H_{bind}$ and *n* value were obtained by fitting with one site binding model. $\Delta G_{bind}$ was calculated based on the relation of $\Delta G_{bind} = -RT\ln K_a = RT\ln K_d$. (-)$T\Delta S_{bind}$ was finally obtained based on the relation of $\Delta G_{bind} = \Delta H_{bind} - T\Delta S_{bind}$.

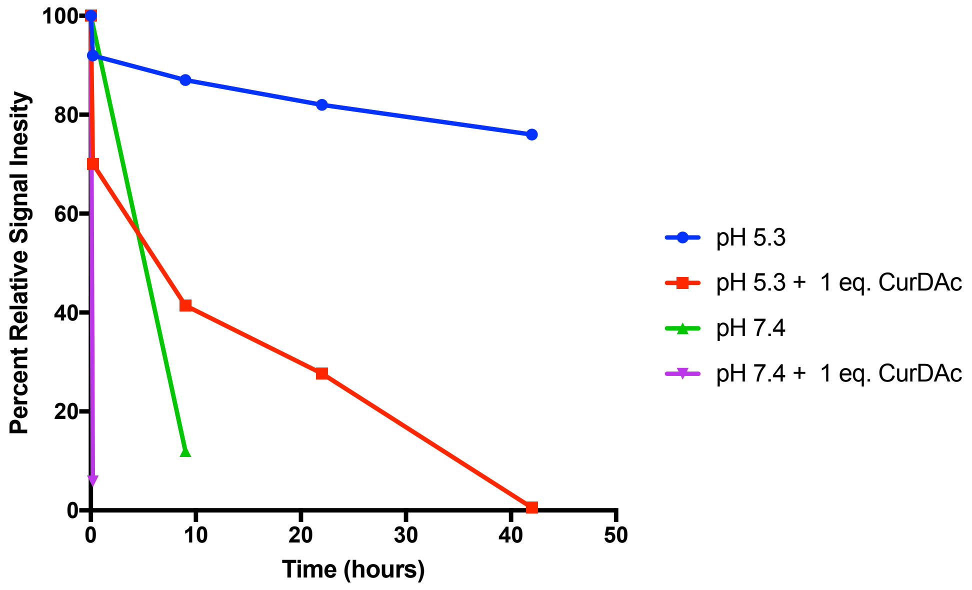


Figure S5. NMR aggregation kinetics of hIAPP and hIAPP + CurDAc as measured by 1D $^{1}H$ signal intensity. 100 μM hIAPP dissolved into pH 7.4 phosphate or pH 5.3 acetate buffers with and without CurDAc were used to measure the proton NMR spectra of the soluble hIAPP peptide. Experimentally measured proton NMR signal intensities are plotted over the time. First spectra were taken immediately after addition of CurDAc to hIAPP monomer and subsequent spectra were taken at various time points.

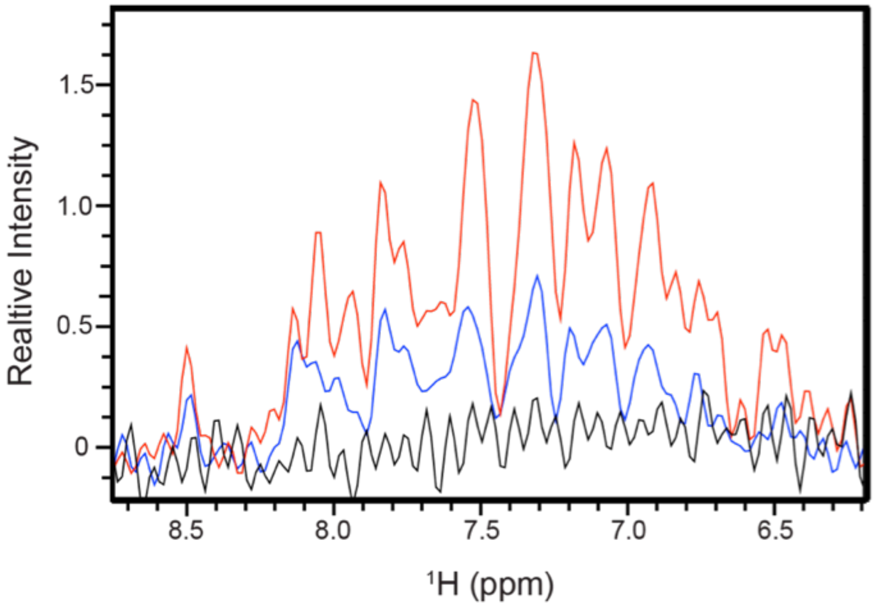


Figure S6. $^{1}$H NMR spectra of hIAPP monomer with 1 equivalent of CurDAc over time obtained from a 850 MHz NMR spectrometer at 25 °C. Red trace spectrum shows 80 μM hIAPP + CurDAc at pH 5.3 acetate buffer at 0 hours. Blue trace spectrum shows the same sample after 9 hours of incubation, and black is after 40 hours.

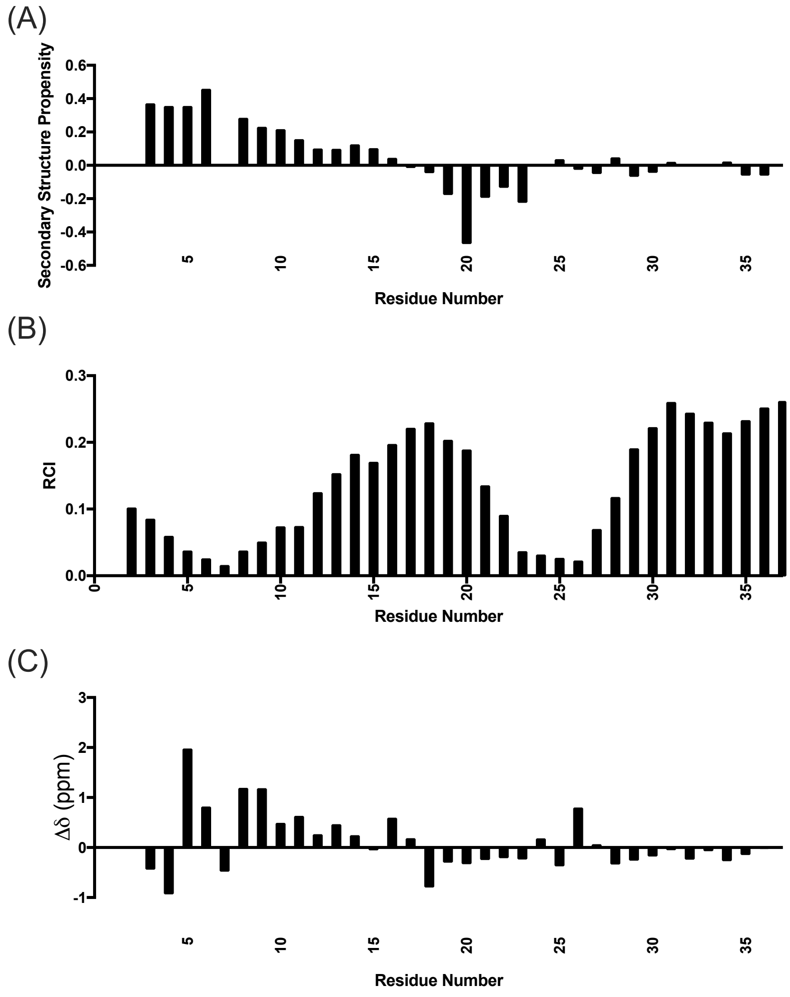


Figure S7. Secondary structure predictions of hIAPP monomer in the presence of CurDAc using the $C_\alpha$, CO, HN chemical shifts. (A) Secondary structure propensities from the Forman Kay group. (B) Random Coil Index from the Wishart group and (C) the $\Delta\delta$ of the $C_\alpha$ chemical shifts. All three show a tendency of an alpha-helical structure at the N-terminus, a small portion of a beta-sheet confirmation near the amyloidogenic core, and then a random coil for the rest of the peptide, which is closely similar to the hIAPP monomer structure.

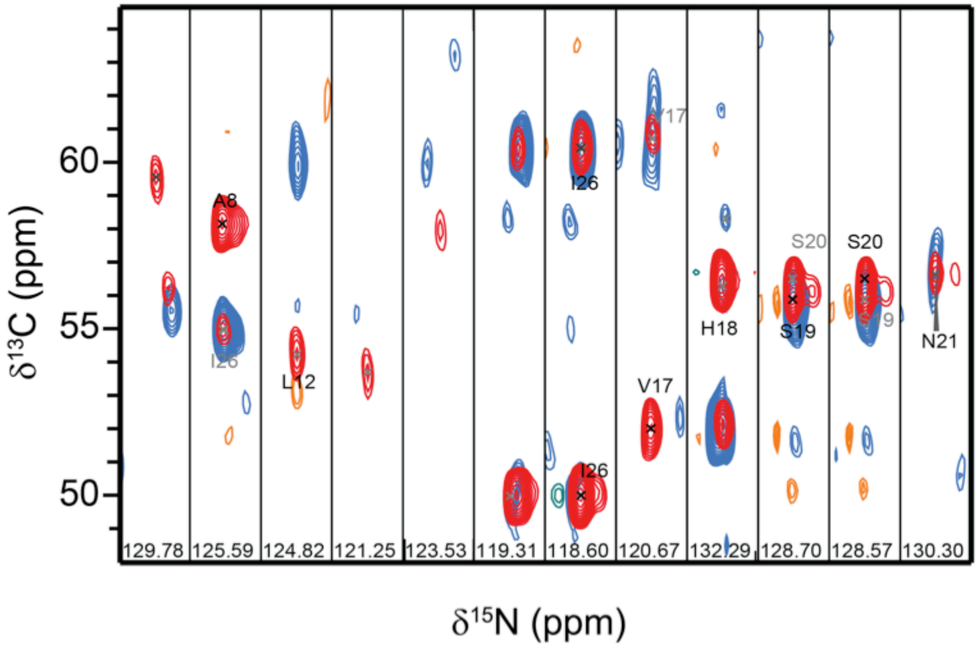


Figure S8. 3D HNCA and HNCOCA experiments performed after 72-hour incubation of hIAPP with CurDAc at a 1:1 molar ratio at pH 7.4 phosphate buffer obtained from a 850 MHz NMR spectrometer at 25 °C.

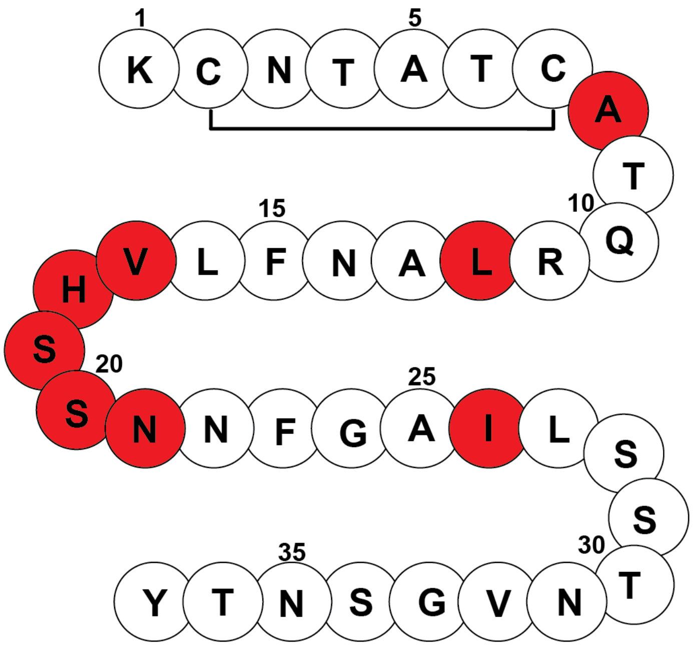


Figure S9. Amino acid sequence model of hIAPP with highlighted residues identified after CurDAc disaggregation in red.

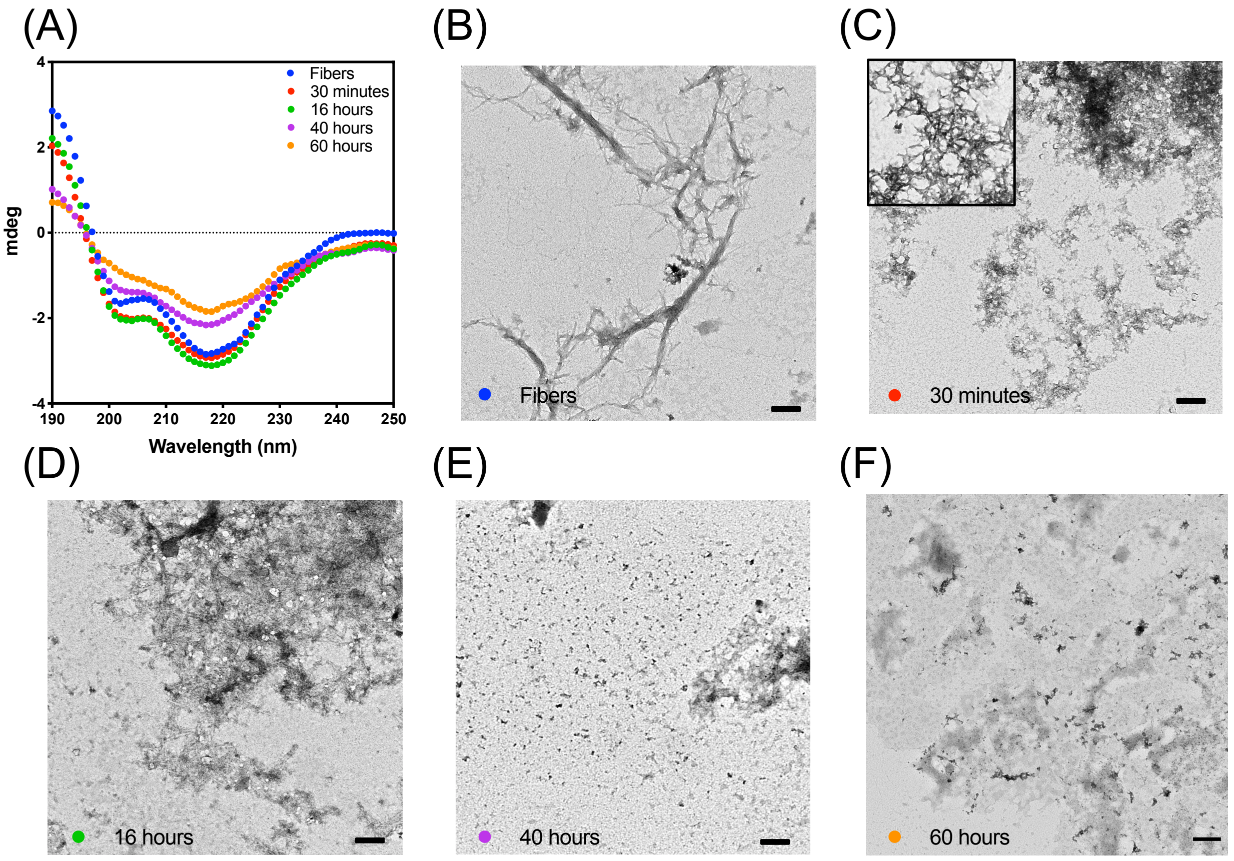


Figure S10. Circular dichroism (CD) and TEM of hIAPP fibers before and after CurDAc treatment. (A) CD spectra of hIAPP fibers in the presence of one molar equivalent of CurDAc over a course of 60 hours in phosphate buffer at pH 7.4. TEM images of hIAPP fibers (B) followed by hIAPP samples in the presence of CurDAc at 30 minutes to 60 hours (C-F) scale bar is 200 nm. Full images are represented in this figure while panels of B, C and F are show in main text figure 3F.

| Sample | Helix 1 | Helix 2 | Total Helix | Anti 1 | Anti 2 | Anti 3 | Para | Turn | Total β-Sheet | Others | NRMSD |
|---|---|---|---|---|---|---|---|---|---|---|---|
| **hIAPP Fiber** | 0 | 0 | 0 | 7.1 | 7.46 | 19.93 | 9.32 | 10.23 | 54.04 | 45.97 | 0.0215 |
| **30 Minutes** | 9.14 | 0 | 9.14 | 0 | 0 | 19.46 | 20.12 | 8.66 | 48.24 | 42.63 | 0.0334 |
| **16 Hours** | 10.49 | 1.83 | 12.32 | 0 | 0 | 16.6 | 19.67 | 7.48 | 43.75 | 43.93 | 0.0366 |
| **40 Hours** | 0 | 2.69 | 2.69 | 0 | 0 | 14.23 | 23.63 | 13.73 | 51.59 | 45.72 | 0.0529 |
| **60 Hours** | 0 | 0 | 0 | 0 | 1.87 | 10.25 | 21.36 | 15.85 | 49.33 | 50.68 | 0.0535 |

Table S2. Parameters obtained by fitting circular dichroism spectra of hIAPP time course disaggregation with CurDAc using BeStSEL. Values are shown in percentage of total content.

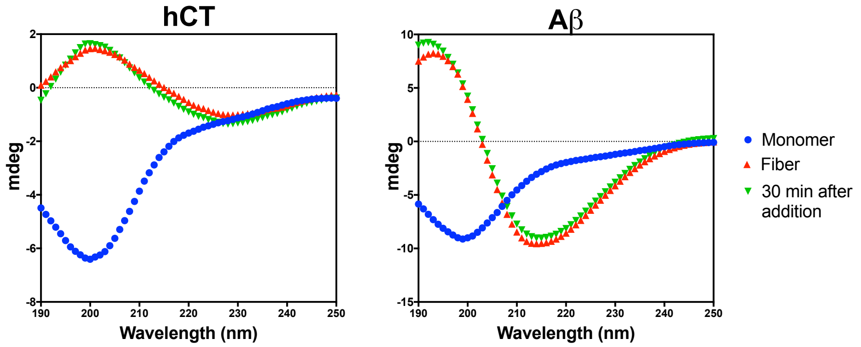


Figure S11. CD spectra of hCT and Aβ before and after treatment with CurDAc. 40 μM peptide monomers in pH 7.4 phosphate buffer show random coil at time zero, and after 24 hours peptides show beta-sheet conformation. hCT was treated with 2 equivalents of CurDAc, and Aβ with 10 equivalents, and then measured again after 30 minutes of incubation.

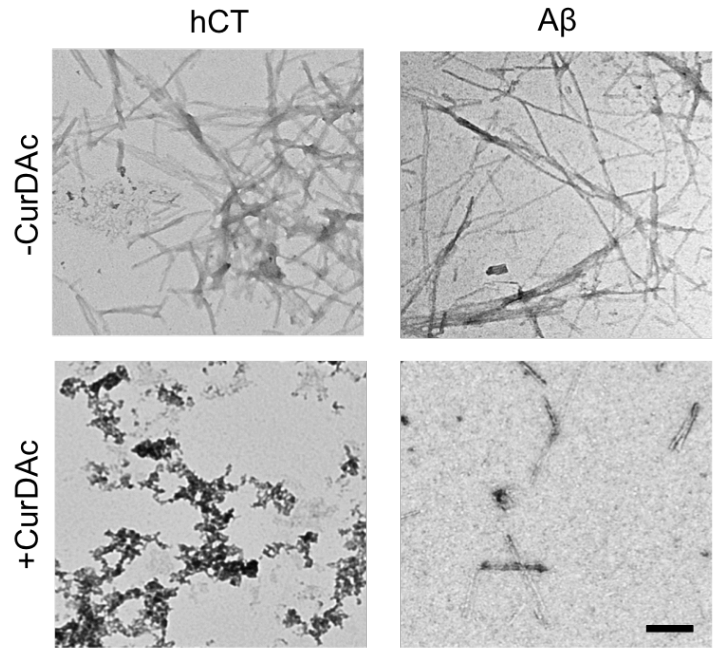


Figure S12. hCT and Aβ TEM images before and after treatment with CurDAc. Top, 40 μM hCT (left) and Aβ (right) after 24 hours shaking at pH 7.4 phosphate buffer at 25°C and 37°C. Bottom, 30 minutes after the addition of CurDAc. 2 equivalents of CurDAc were added to hCT and 10 Equivalents were added to Aβ. Scale bar is 100 nm.

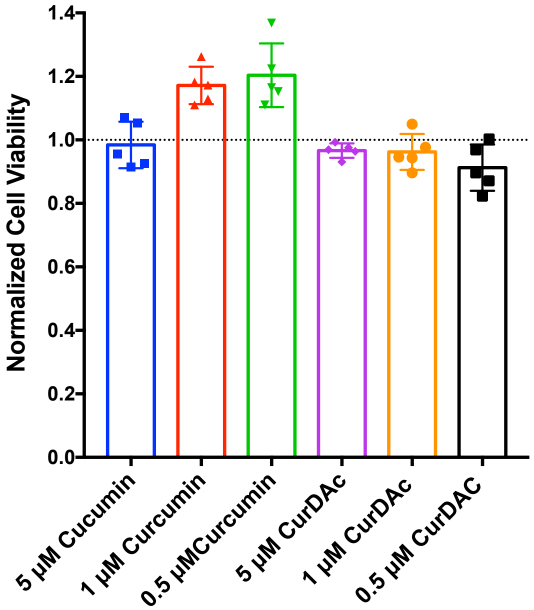


Figure S13. MTT cell toxicity assay of curcumin and CurDAc. RIN-5F cells treated with varying amounts of curcumin or CurDAc. MTT reduction was measured after 24 hours.

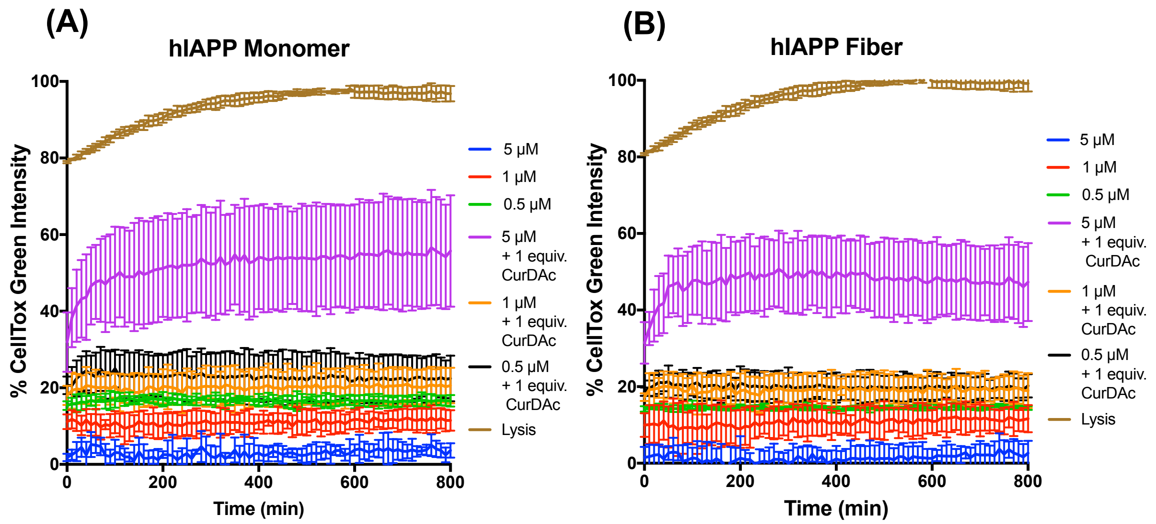


Figure S14. RIN-5F cell death monitored every 10 minutes for 13 hours by CellTox Green fluorescence in the presence of hIAPP monomers or fibers with varying amounts of CurDAc.

| Samples Compared | Significant? | Summary | Adjusted P Value |
|---|---|---|---|
| hIAPP monomer MTT Experiments | | | |
| 5 μM vs. 5 μM + 1 equiv. CurDAc | Yes | **** | <0.0001 |
| 1 μM vs. 1 μM + 1 equiv. CurDAc | Yes | **** | <0.0001 |
| 0.5 μM vs. 0.5 μM + 1 equiv. CurDAc | Yes | **** | <0.0001 |
| 5 μM + 1 equiv. CurDAc vs. 1 μM + 1 equiv. CurDAc | Yes | ** | 0.0034 |
| 5 μM + 1 equiv. CurDAc vs. 0.5 μM + 1 equiv. CurDAc | Yes | **** | <0.0001 |
| 1 μM + 1 equiv. CurDAc vs. 0.5 μM + 1 equiv. CurDAc | No | ns | 0.0855 |
| hIAPP Fiber MTT Experiments | | | |
| 5 μM vs. 5 μM + 1 equiv. CurDAc | No | ns | 0.9985 |
| 1 μM vs. 1 μM + 1 equiv. CurDAc | Yes | **** | <0.0001 |
| 0.5 μM vs. 0.5 μM + 1 equiv. CurDAc | Yes | **** | <0.0001 |
| 5 μM + 1 equiv. CurDAc vs. 1 μM + 1 equiv. CurDAc | Yes | **** | <0.0001 |
| 5 μM + 1 equiv. CurDAc vs. 0.5 μM + 1 equiv. CurDAc | Yes | **** | <0.0001 |
| 1 μM + 1 equiv. CurDAc vs. 0.5 μM + 1 equiv. CurDAc | No | ns | 0.1464 |
| hIAPP Monomer CellTox Green Experiments |  |  |  |
| 5 μM vs. 5 μM + 1 equiv. CurDAc | Yes | **** | <0.0001 |
| 1 μM vs. 1 μM + 1 equiv. CurDAc | No | ns | 0.5995 |
| 0.5 μM vs. 0.5 μM + 1 equiv. CurDAc | No | ns | 0.9128 |
| 5 μM + 1 equiv. CurDAc vs. 1 μM + 1 equiv. CurDAc | Yes | *** | 0.0002 |
| 5 μM + 1 equiv. CurDAc vs. 0.5 μM + 1 equiv. CurDAc | Yes | *** | 0.0003 |
| 1 μM + 1 equiv. CurDAc vs. 0.5 μM + 1 equiv. CurDAc | No | ns | 0.9998 |
| hIAPP Fiber CellTox Green Experiments | | | |
| 5 μM vs. 5 μM + 1 equiv. CurDAc | Yes | **** | <0.0001 |
| 1 μM vs. 1 μM + 1 equiv. CurDAc | No | ns | 0.5284 |
| 0.5 μM vs. 0.5 μM + 1 equiv. CurDAc | No | ns | 0.7522 |
| 5 μM + 1 equiv. CurDAc vs. 1 μM + 1 equiv. CurDAc | Yes | **** | <0.0001 |
| 5 μM + 1 equiv. CurDAc vs. 0.5 μM + 1 equiv. CurDAc | Yes | *** | 0.0001 |
| 1 μM + 1 equiv. CurDAc vs. 0.5 μM + 1 equiv. CurDAc | No | ns | 0.9999 |

Table S3. Statistical significance calculated by one-way ANOVA using Tukey's multiple comparison tests. Comparisons were done between cells treated with hIAPP with and without CurDAc.
